# A multimodal framework for detecting direct and indirect gene-gene interactions from large expression compendium

**DOI:** 10.1101/680116

**Authors:** Lu Zhang, Jia Xing Chen, Shuai Cheng Li

## Abstract

The fast accumulation of high-throughput gene expression data provides us an unprecedented opportunity to understand the gene interactions and prioritize disease candidate genes. However, these data are typically noisy and highly heterogeneous, complicating their use in constructing large expression compendium. Recent studies suggest that the collective expression pattern can be better modeled by Gaussian mixtures. This motivates our present work, which applies a Multimodal framework (MMF) to depict the gene expression profiles. MMF introduces two new statistics: Multimodal Mutual Information and Multimodal Direct Information. Through extensive simulations, MMF outperforms other approaches for detecting gene co-expressions or gene regulatory interactions, regardless of the level of noise or strength of interactions. In the principal component analysis for very large collections of expression data, the use of MMI enables more biologically meaningful spaces to be extracted than the use of Pearson correlation. The practical use of MMF is further demonstrated with three biological applications: 1. Prioritizing *KIF1A* as the candidate causal gene of hereditary spastic paraparesis from familial exome sequencing data; 2. Detecting *ANK2* as the ‘hot genes’ for autism spectrum disorders, derived from exome sequencing family based study; 3. Predicting the microRNA target genes based on both sequence and expression information.

## 1 INTRODUCTION

A massive amount of biological data have been accumulated in the past decades through the widespread application of high-throughput technology in molecular biology. These data contain more valuable knowledge which may be overlooked by inspecting each dataset individually. The gene expression datasets, which are publicly available in several sites (such as Gene Expression Omnibus (GEO) Barrett et al. (2013), ArrayExpress Kolesnikov et al. (2015) and etc.), provide us an unprecedented opportunity to discover valuable information. Different from structuralized data, analysis for gene expression is a data-driven and unsupervised learning task. Integrating different data sources can enable us eliminate the issues caused by poor quality and incomplete training set. The analyses of large amount of gene expression collections have been applied in predicting gene functions Fehrmann et al. (2015), prioritizing disease candidate genes van Dam et al. (2015), predicting microRNA targets Gennarino et al. (2009, 2012), constructing gene co-expression network Lee et al. (2004), *etc*.

These analyses rely heavily on the accurate detection of gene-gene interactions from large expression compendium, several computational approaches have been developed for this purpose. Computational approaches typically define gene-gene interactions by gene co-expression; the rationale is that, when two genes demonstrate correlated expression patterns, they are likely to interact.

Pearson correlation followed by meta-analysis is widely employed to reconstruct gene co-expression network from large expression data. This approach benefits from its straightforward interpretation and computational efficiency. Lee Lee et al. (2004) performs the first study to prove the reproducibility of co-expression relationship across multiple datasets by counting the significant gene correlations from 3,924 microarrays. The co-expressed gene pairs, supported by multiple datasets, are strongly enriched in biological function. MEFIT Huttenhower et al. (2006) utilizes a scalable Bayesian framework to integrate the Z-scores that are transformed from Pearson correlations between gene pairs, and is proved to be superior to the other integration methods. However, Pearson correlation can only capture linear correlation, which is not always the case for gene-gene interactions. This has prompted the use of mutual information (MI) in the inference of gene-gene interactions. The MI of two genes is a measure of their mutual dependence which can detect both linear and nonlinear dependencies Brunel et al. (2010); Meyer et al. (2008); Luo et al. (2008). For gene expression data, MI is computed by discretization through B-spline smoothing Daub et al. (2004), or by assuming Gaussian distributions Margolin et al. (2006).

Although gene co-expression reveals the dependencies between genes, it cannot distinguish between the direct and transitive dependencies–the latter is often considered as false positives for gene regulatory relations as the regulators bind to their target genes physically. Many approaches have been proposed to remedy this. ARACNE Margolin et al. (2006) considers the lowest MI value among any triplet of genes as a transitive edge based on Data Processing Inequality. The CLR Faith et al. (2007) algorithm transforms the MI to Z-score to remove background promiscuous gene correlations. GENIE3 Huynh-Thu et al. (2010) creates a tree-based ensemble model for each target gene to predict and rank the potential regulatory links. TIGRESS Haury et al. (2012) proposes a robust and accurate method for stability selection to improve the feature selection in least angle regression for each target gene. If gene expression data follow a Gaussian distribution, transitive elements in the gene covariance matrix can be eliminated by using precision matrix, as demonstrated by MaxEnt Lezon et al. (2006) based on the maximum entropy principle.

The sample size of the dataset is an important factor affecting the accuracy of inferring the gene-gene interactions, as several studies indicated before. One approach to alleviate this issue is to integrate expression data from available databases. HOCTAR Gennarino et al. (2009) infers the microRNA targets by considering the expression correlations between microRNA host gene and its potential target genes, which are calculated by integrating 3,445 different microarray hybridization experiments from Affymetrix HGU133A. However, HOCTAR neglects the transitive effects, which may report the genes that co-express with the real targets rather than to be regulated by the microRNA. Though integrating multiple datasets increases the sample size, the data will be severely obstructed by the heterogeneity: the collected samples are produced from different tissues, by different platforms, by different RNA extraction methods, and with varying qualities.

Methods such as Pearson correlation followed by meta-analysis can relief the heterogeneous issue across datasets, but still are inane to the inner heterogeneity within each dataset. We note a related study with the gene expressions of tumor tissues, which are often confounded by their surrounding normal tissues or mixed with different subclones Navin et al. (2011). To distinguish these tissues, TEMT Li and Xie (2013) models gene expression with Gaussian mixture models. We also note a result by Kim Kim et al. (2010) over the large expression compendium, where it found the majority of gene expressions follow multimodal distributions: 48.9% and 34.7% of probes should be modeled as for bi- and tri-modes distributions. Motivated by these results, we use Gaussian mixtures on our framework to infer gene interactions.

Previous methods model the gene expressions according to their “global features”–following a common distribution. But actual situation is not always the case. Some genes merely express in a specific cellular condition or tissue Wang et al. (2014), they may be modeled by appropriate “local features”–following the combination of multiple distributions. To model the “local features”, we assume that the gene expressions are sampled from different distributions rather than independent and identically distributed random variables. Hence, we propose a Multimodal Framework (MMF) that depicts the large gene expression data explicitly by Gaussian mixture models. Under this framework, the correlations are evaluated more accurately through a new measure—Multimodal Mutual Information (MMI). MMF also allows a new measure called Multimodal Direct Information (MDI) to identify regulation relationship free from the influence of transitive correlations. These two measures form the basis of our framework to identify gene interactions from integrated large expression datasets.

When comparing to the other methods for inferring gene-gene interactions, MMI and MDI demonstrate superior accuracy and noise tolerance according to the simulation results. We further successfully apply MMI and MDI to three biomedical problems and obtain encouraging results: 1. MMI identifies *KIF1A* as the causal gene of hereditary spastic paraparesis (HSP) correctly from familial exome sequencing data by detecting the strongest co-expression with the established disease casual genes; 2. MDI identifies *ANK2* as the ‘hot gene’ from exome sequencing familial study for autism spectrum disorders (ASDs); 3. MDI predicts the targets of microRNAs transcribed from the intragenic regions accurately. These experiments demonstrate the effectiveness of MMF in identifying gene interactions from large gene expression compendium.

## 2 MATERIALS AND METHODS

### 2.1 Multimodal framework

The MMF is specifically designed for calculating gene-gene interactions from noisy and heterogeneous expression datasets. MMF considers the “local feature” under the assumption that the expression data of each gene are sampled from Gaussian mixture models rather than one Gaussian distribution (**Materials and methods**). MMI is first proposed to evaluate the co-expression between gene pairs based on Gaussian mixtures. We further implement MDI to eliminate transitive interactions according to maximum entropy principle (**Materials and methods** and **Supplementary note**), which shed light on the identification of regulatory interactions and master regulators. The key innovation of MMF is considering both “outer” and “inner” local features. The “outer” local feature refers to the local probability of expression profiles from the same mode; the “inner” local feature is the correlation of gene expressions in the same mode. Under the framework, we define a measure called MMI to capture the gene co-expression, and another called MDI to capture gene regulatory interactions. The entire framework of MMF is showed in Figure fig:Figure1.

**Figure 1.**
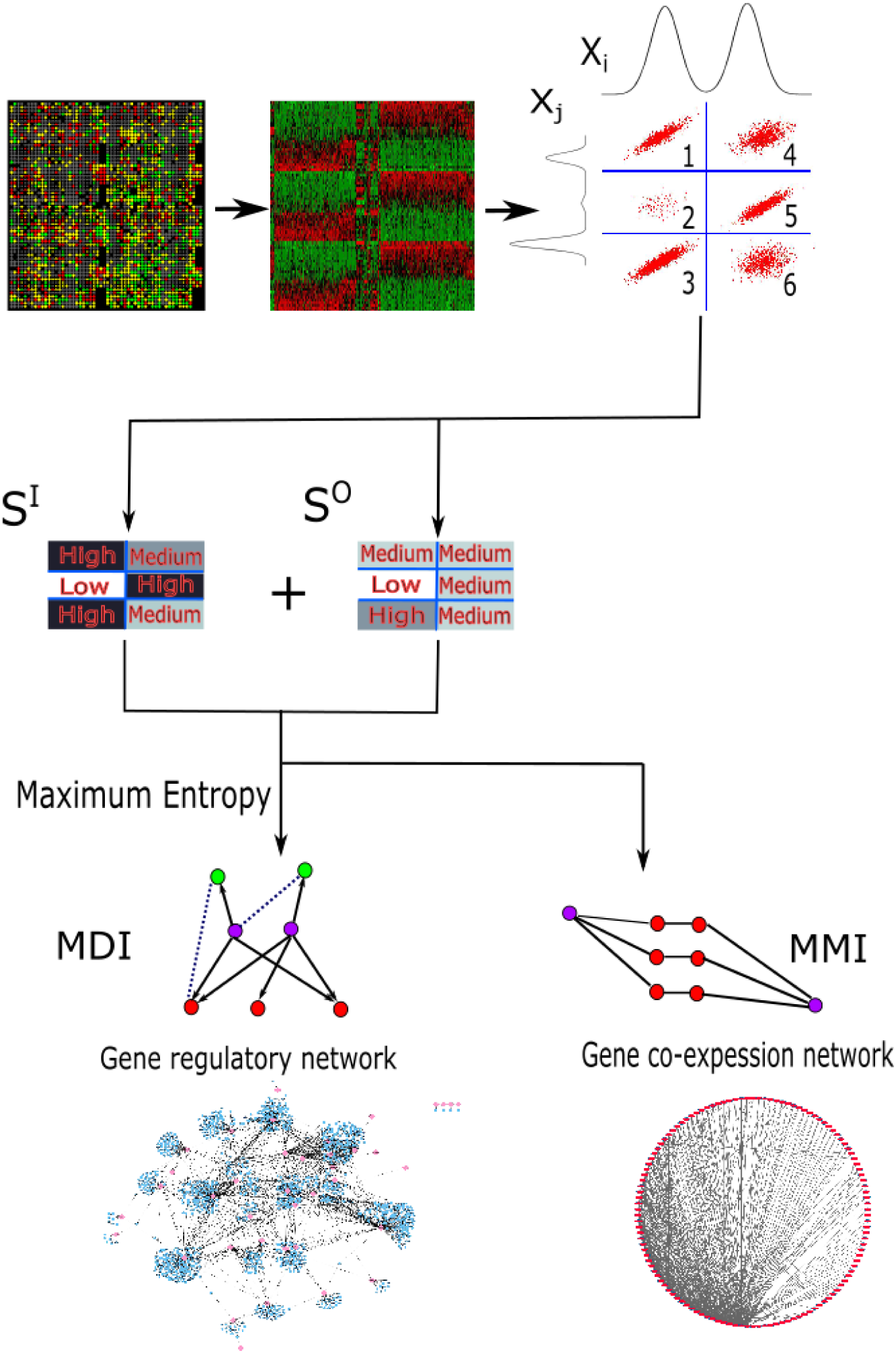
The procedure and purpose for MMI and MDI under Multimodal framework. Two genes *X*_*i*_ and *X*_*j*_ come from Gaussian Mixture models with two and three modes, respectively. The samples are divided into six bins as the Cartesian product of the clusters for *X*_*i*_ and *X*_*j*_. The expression profiles are highly co-expressed in the 1st, 3rd and 5th bins. The 4th and 6th bins are marginally correlated. There are only a few samples in the 2nd bin with weak correlation. The *S*^*I*^ and *S*^*O*^ are calculated with deeper color demonstrating stronger covariance. MMI calculates the co-expression between gene pairs (purple circles), regardless whether there are transitive nodes (red circles) between them. MDI captures the regulatory interactions (arrows present regulation directions), the transitive interactions are eliminated (dashed lines).

### 2.2 Gene expression data integration

We collect gene expression profiles from GEO and choose the most comprehensive array platform HG U133 Plus 2.0 (**TableS1**); All of the samples are processed together to examine the performance of MMF in the global network. Poor quality chips are removed through the `affyQCReport` package from Bioconductor. Likewise, `GCRMA` is utilized to extract the log scale expression profiles for each probe followed by quantile normalization. The intensity of probe smaller than two is regarded as missing value and imputed by `impute` package from Bioconductor. Some genes are annotated by multiple probes, their expression profiles are computed by averaging those probe expressions.

### 2.3 Determine the number of modes for each gene

We assume that the integrated expression data is an *m* × *n* matrix, *D* = (*d*_*i,j*_)_*m*×*n*_, where each row *i* (denoted *D*_*i,•*_) represents a sample, and each column *j* (denoted *D*_*•,j*_) represents the expression profiles of one gene across all the samples.

We now describe how MMF models the underlying distribution that gives rise to this expression data. First, we group the expression profiles of each gene into clusters, where each cluster is assumed to form a Gaussian distribution. The expression profiles of any gene pair are partitioned according to the Cartesian product of the clusters from the respective genes. Each partition follows bivariate Gaussian distribution. Denote the clusters of gene *j* as 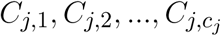, where *c*_*j*_ is the number of clusters for gene *j*. The clusters for each gene is determined by maximizing the total log-likelihood of the Gaussian distributions formed. The log-likelihood of expression profiles 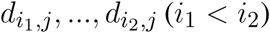 to construct a Gaussian distribution is computed as

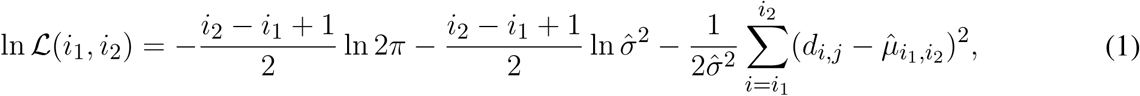

where 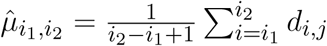 and 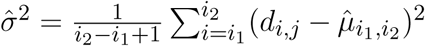. Our aim is to partition the data, *d*_1*,j*_*, …, d*_*m,j*_, into distributions such that the sum of log-likelihoods from all the distributions is maximized. This can be solved using dynamic programming as follows.

Without loss of generality, we assume the expression profiles of gene *j* are sorted; that is, *d*_1*,j*_ ≤ *d*_2*,j*_ ≤ *…* ≤ *d*_*m,j*_ (it is clear that such a sorting can be performed very efficiently).

Let *T*(*i, k*) denote the maximum likelihood by clustering the data *d*_1*,j*_ ≤ *d*_2*,j*_ ≤ *…* ≤*d*_*i,j*_ into *k* clusters. Then, the following recurrence relations can be formulated,

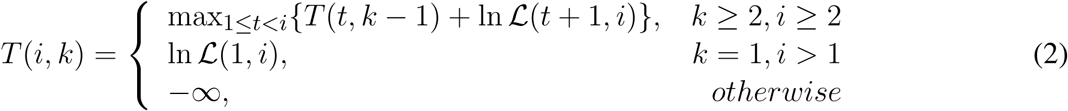

Hence, the maximum *T* (*i, k*) and its corresponding clusters can be calculated through dynamic programming for a given *k*.

### 2.4 Models for expression profiles of a single gene

We assume a random variable *X*_*j*_ for the expression of gene *j*. Following our framework, *X*_*j*_ is decomposed into *c*_*j*_ Gaussian random variables, denoted 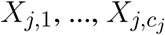. Denote the density function for cluster *C*_*j,k*_ as *gX*_*j,k*_ (*x*), 1≤ *k* ≤ *c*_*j*_. Then, the expression profiles for gene *j* is distributed according to the density function

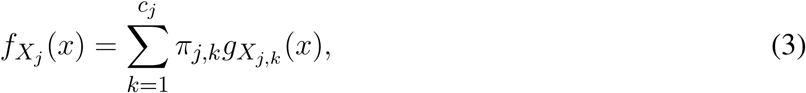

where

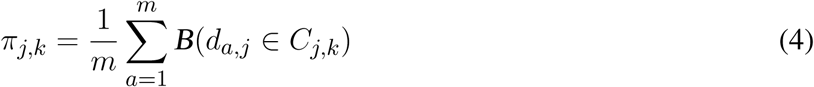

is the proportion of samples in cluster *C*_*j,k*_; here, *B* denotes Boolean function.

It is possible to introduce a notation of entropy for each gene whose expression profiles follow Gaussian mixture models:

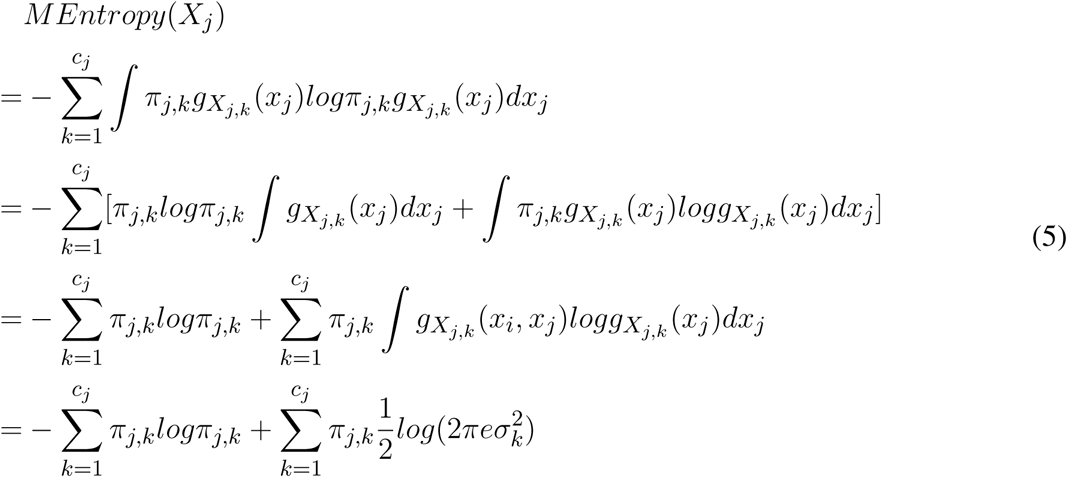

in which, *σ*_*k*_ denotes the standard deviation for the *k*th Gaussian distribution. *MEntropy* is used to normalize MMI and MDI to [0, 1] in Section Sec:backgroundnoise.

### 2.5 Multimodal mutual information

Computing MMI consists of four major steps: first, the expression profiles for each gene are clustered by assuming that each cluster is Gaussianly distributed; second, “outer” MI is computed by aggregating the Kullback-Leibler divergence from the discretized gene expression profiles; third, “inner” MI is calculated for each cluster formed by any two genes; fourth, the MMI of two genes is calculated by aggregating the “outer” and “inner” MIs across all the associated clusters.

### 2.6 Models for expression profiles of gene pairs

We capture the relations between expression profiles of two genes by bivariate Gaussian mixture models. Given the expression profiles of gene *i* and *j* (1 ≤ *i, j* ≤ *n*), we partition the data into *c*_*i*_ × *c*_*j*_ bins; that is, we take the Cartesian product of the clusters for gene pair *i* and *j*. We model each bin as a bivariate Gaussian distribution, and denote the density function of each distribution as 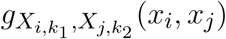, where 1 ≤ *k*_1_ ≤ *c*_*i*_ and 1 ≤ *k*_2_ ≤ *c*_*j*_. The expressions of genes *i* and *j* is a mixture models with joint density function

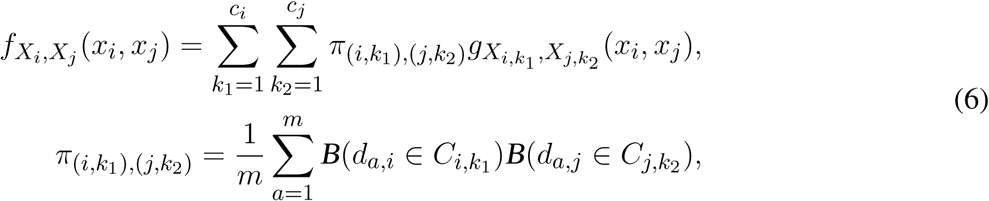

where 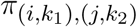 is the proportion of samples shared by cluster 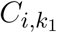 and 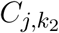. We assume that the marginal distributions of 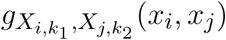 are *gX* _*i,k*1_ (*x*_*i*_) and *gX*_*j,k*2_ (*x*_*j*_). Hence, the only parameter left to be estimated is the covariance matrix (correlation matrix). Denote the covariance between variable 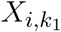 and 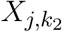 as 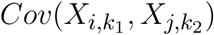. Notice that we cannot utilize the covariance of shared expression profiles between 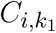 and 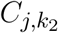, we need to guarantee the marginal distributions of each bin are invariant.

### 2.7 Covariance matrix estimation

We calculate the covariance matrix *S*^*I*^ to capture the covariance in each bin, whose entry 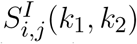 denotes the covariance between two variables 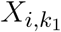 and 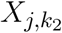. We first construct 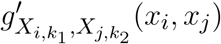 that according to the expression profiles shared between 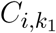 and 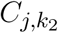. Assuming that 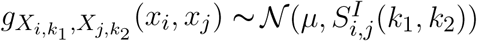 and 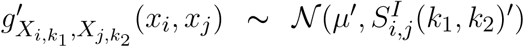, we can calculate 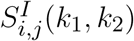 by minimizing these Kullback-Leibler divergence between the two distributions by Eq. eqn:opt:

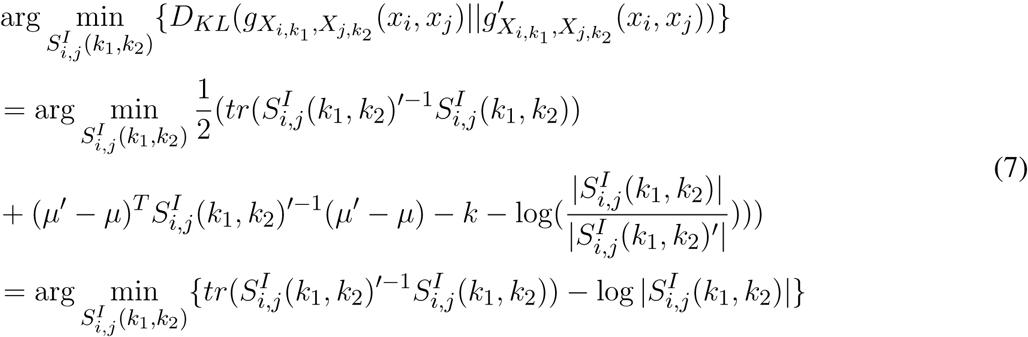

### 2.8 Aggregating the “outer” and “inner” mutual information

After calculating the mixture distributions and their parameters, we need to aggregate MI from each bin to detect the interactions between two genes. By assuming that *X*_*i,k*_ follows a Gaussian distribution, the mutual information between *X*_*i*_ and *X*_*j*_ is estimated as

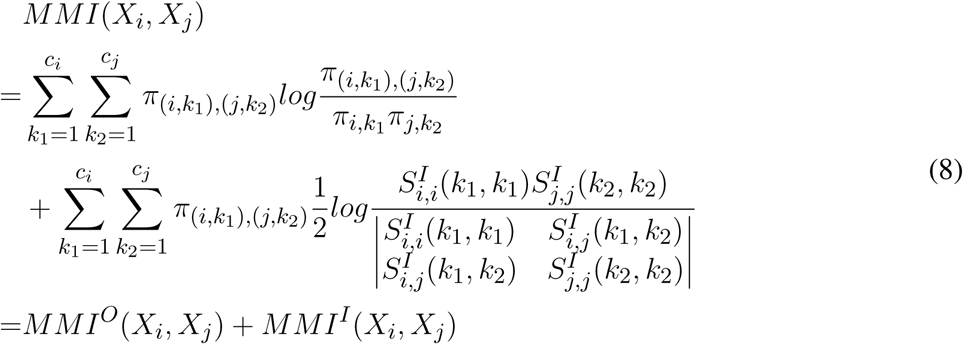

in which | • | denotes matrix determinant. From Eq. eqn:MMI, we observe that MMI is calculated by aggregating two types of mutual information: *MMI*^*O*^(*X*_*i*_*, X*_*j*_), which we refer to as “outer” mutual information, and *MMI*^*I*^(*X*_*i*_*, X*_*j*_), which we refer to “inner” mutual information. The “outer” mutual information is calculated by discretizing the continuous expression profiles into small bins, and is basically the same as the MI calculated for relevance networks (Butte and Kohane, 2000). The “inner” mutual information is the weighted aggregation of mutual information for each bin.

### 2.9 Multimodal Direct Information

To remove transitive interactions between any gene pairs, we introduce a measure—Multimodal Direct Information— which is enhanced from MMI based on maximum entropy principle. The “outer” part of MDI, *MDI*^*O*^, is modified from the direct-coupling analysis (DCA) (Morcos, Pagnani, Lunt, Bertolino, Marks, Sander et al., 2011), that identifies the co-evolution between protein residuals. The inner part of MDI, *MDI*^*I*^, is similar to *MMI*^*I*^, but with the covariance matrix *S*^*I*^ exchanged with a precision matrix, while ensuring the marginal distributions are invariant.

### 2.10 “Outer” MDI

DCA has been successfully applied to identify co-evolved protein residuals by removing false transitive connections. *MDI*^*O*^ is based on a similar technique as DCA, but modified for gene expression data. First, *MDI*^*O*^ introduces *pseudosamples λ* in each bin to avoid insufficient samples. Let *pseudosamples* be uniformly distributed across all the bins. Then, *π*_*j,k*_ and 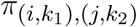 are rewritten as:

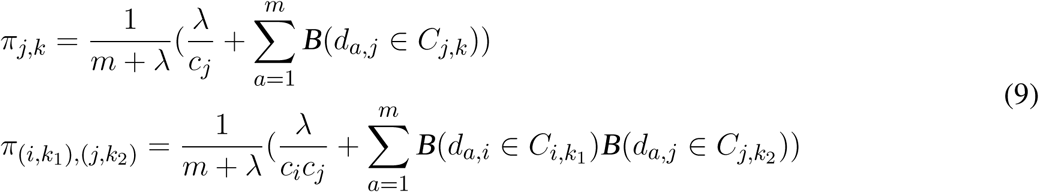

the covariance between any pair of genes (*i, j*) for the bin (*k*_1_*, k*_2_) in *MDI*^*O*^ is given by

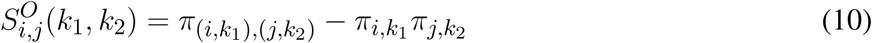

### 2.11 “Inner” MDI

Rather than normalizing each term as in *MMI*^*I*^ by the number of samples grouped in the particular bin, *MDI*^*I*^ introduces an *“inner” psuedocount* to provide the clusters for each gene the same sample size. For the *k*th bin of gene *j*, we denote the average expression of the samples in *C*_*j,k*_ is 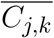. For each sample which is not a member of the cluster 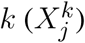, its value is replaced with 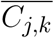 in *MDI*^*I*^. The covariance for the bin *k*_1_*, k*_2_ of gene *i, j* is

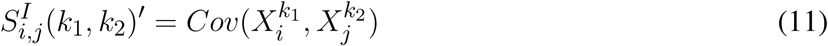

The same as MMI, 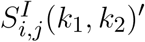 is further transformed to 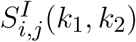 according to Eq.eqn:opt.

#### Precision matrix

In order to calculate the precision matrix more efficiently and accurately, we introduce a regularization parameter *η*. The precision matrix Θ is calculated as

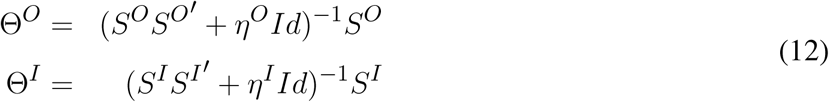

### 2.12 Aggregate “outer” and “inner” Direct information

Finally, MDI is calculated by aggregating the *MDI*^*O*^(*i, j*) and *MDI*^*I*^(*i, j*) across all the bins.

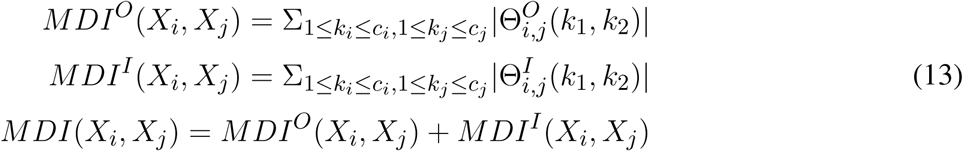

### 2.13 Background noise elimination

MMI and MDI can be further adjusted to eliminate the background influence, or noise (Eq.eqn:backgroundnoise). After that, we rescale MMI and MDI to [0, 1] by dividing them with their upperbound max*{MEntropy*(*X*_*i*_)*, MEntropy*(*X*_*j*_)*}* ((Eq.eqn:maximum)). In this paper, we perform this step by default, and simply write *MMI*_*adj*_ and *MDI*_*adj*_ as *MMI* and *MDI*.

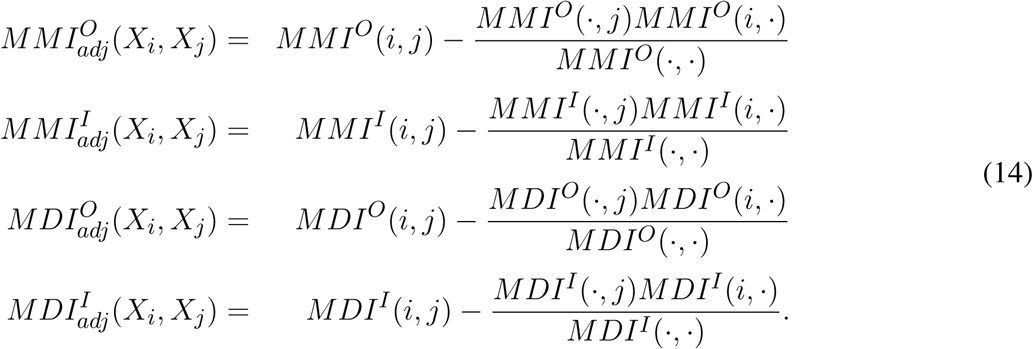

and

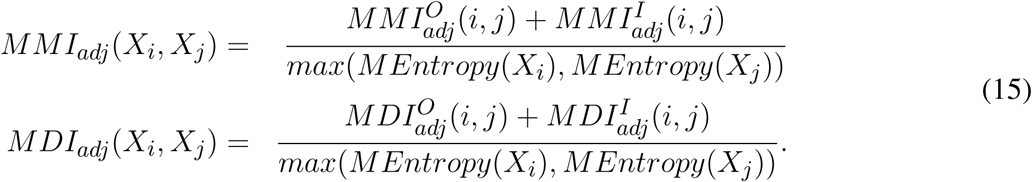

## 3 RESULTS

### 3.1 Simulation for evaluating MMI and MDI

We follow closely the procedures in SIMON and TIBSHIRANI (2012) for simulation. We sample the gene pair expression profiles from bivariate Gaussian mixture distributions (with two modes) with the covariance changing from 0.1 to 0.9. The empirical distribution is constructed by the same procedure, implying the same distribution but with randomly assigned covariances.

We evaluate eight measures besides MMI: 1. Pearson correlation; 2. Spearman correlation; 3. Kendall’s rank correlation; 4. Mutual information based on kernel density estimation (MI(KDE)); 5. Mutual information based on B-spline (MI(bspline), with 10 partitions for each gene); 6. Maximal information coefficient (MIC); 7. Maximum Asymmetry Score (MAS); 8. Maximum Edge Value (MEV). These measures are evaluated by their power, that is, the proportion of simulations exceed the top 5% correlations from the empirical distributions. To evaluate the noise tolerance of these correlation measures, we introduce uniform distributed noises weighted by 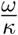 to the simulated expression profiles, where “*ω*” presents the amount of noise and “*κ*” denotes the noise level. We assign “*ω*” to a constant value 3 and set “*κ*” to change from 0 to 3 in the simulations. We also evaluate MMI with different cluster numbers to assess its sensitivity to clustering error.

We test MDI against five famous approaches for inferring regulatory interactions: ARACNE, CLR, GENIE3, MaxEnt and TIGRESS. In order to explicitly reflect the nature of direct interaction, we construct a tree structure, in which each node has only one parent (except the root) and merely directly interacts with its parent. In other words, the expression profiles of offsprings are totally determined by their parents. The expression profile for each node is sampled from a Gaussian mixture models with two modes. The joint distribution of a parent and one of its offsprings is a bivariate Gaussian distribution with specified covariance. The area under ROC (AUC) is applied to evaluate the methods’ performances. The involved noise signals are the same as those applied for the simulations for MMI.

### 3.2 Simulation results for MMI

We perform 1000 simulations to construct empirical distributions. MMI obtains higher powers than the other measures in all the simulations (Figure fig:Figure2), regardless of the magnitude of covariance or noise signals exist. MMI explicitly groups the samples for each gene into two modes followed by aggregating four bins together, where the correlation for each bin is calculated independently. This partitioning strategy makes the expression profiles for each bin follow a Gaussian distribution, which makes MMI easier to capture their correlations.

**Figure 2.**
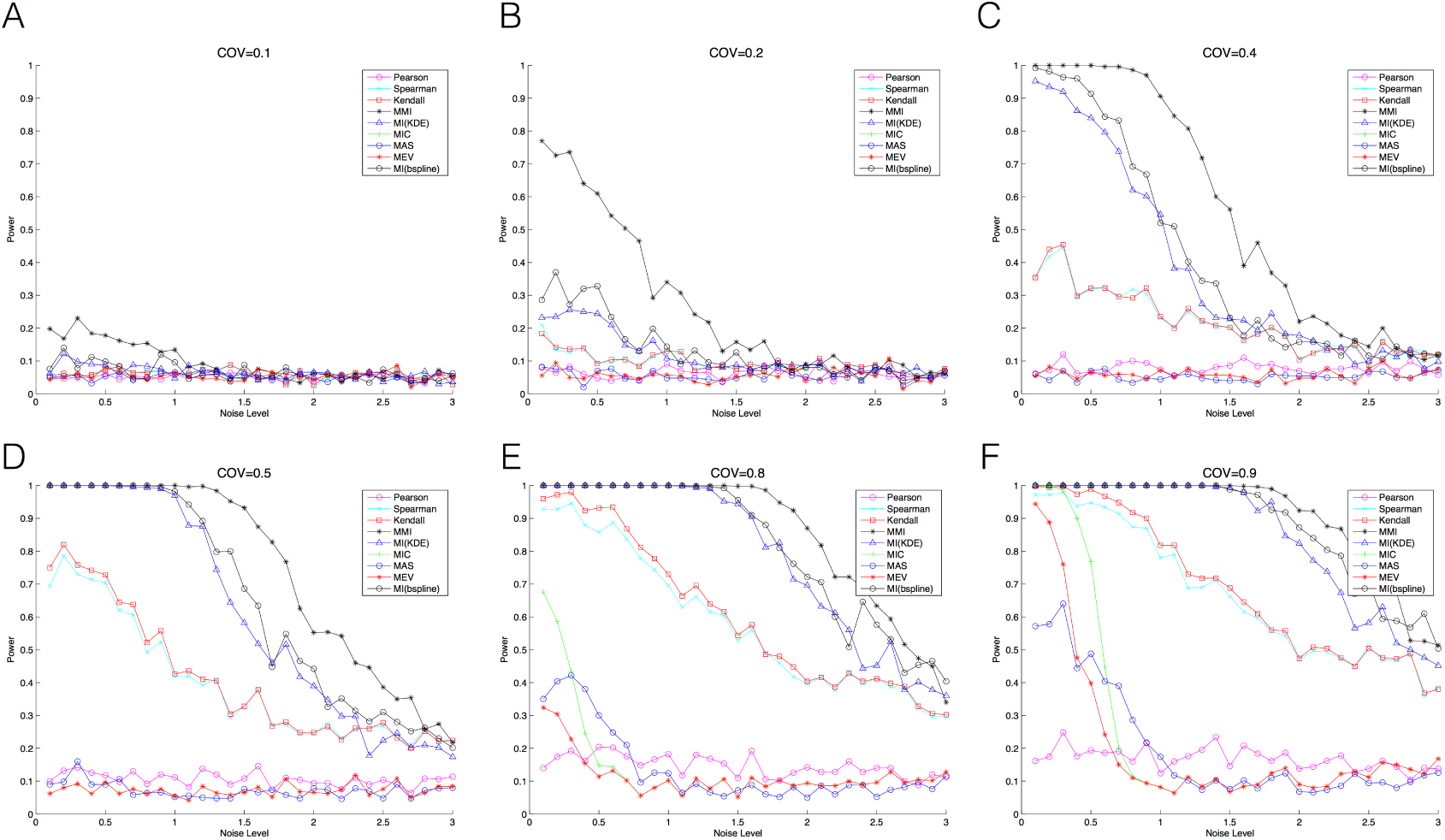
Power and noise tolerance comparing MMI with other methods in simulation data. Simulation Data are sampled from bivariate Gaussian mixture models with different covariances. We add different amount of noise in the simulation data. **(A)** Covariance=0.1, **(B)** Covariance=0.2, **(C)** Covariance=0.4, **(D)** Covariance=0.5, **(E)** Covariance=0.8, **(F)** Covariance=0.9.

It is critical to lessen the influence of noise that may introduce the false positive results. By considering the “local feature”, MMI has better noise tolerance than other measures for uniform distributed noise. The noise is not always uniformly distributed across different studies or platforms. When heavy noise affects only a particular proportion of expression profiles, MMI tolerates the noises as they are usually grouped into isolate bins as the second bin illustrated in Figure fig:Figure1.

The second best method is MI(bspline), a non-parametric method which approximates the probabilistic density function by discretizing the continuous expressions into bins. While MI(bspline) seems to capture the information of *MMI*^*O*^, but neglects the correlation in each bin. The other MI estimation, MI(KDE), utilizes Gaussian kernel to approximate the entire distribution to a Gaussian mixture models. The measure is demonstrably robust, but has an issue with efficiency–making it difficult to be applied to large expression compendium. Two rank based statistics, Spearman’s rank correlation coefficient and Kendall’s rank correlation, perform acceptable for high variance, but are found lacking when the covariance is small.

Considering MMI’s sensitivity to errors in terms of the cluster number for each gene, we assign different cluster number, from 2 to 5, on a Gaussian mixtures of two modes. We denote these MMI instances as MMI(2), MMI(3), MMI(4) and MMI(5). The results are as demonstrated in Figure fig:Figure3. As expected, MMI(2) performs the best in all cases. The performance deteriorates as the cluster number deviates further from the true value. This indicates that the correctness in the number of cluster is crucial to MMI’s performance. However, we note even the performance of MI(5) is comparable to the other measures’.

**Figure 3.**
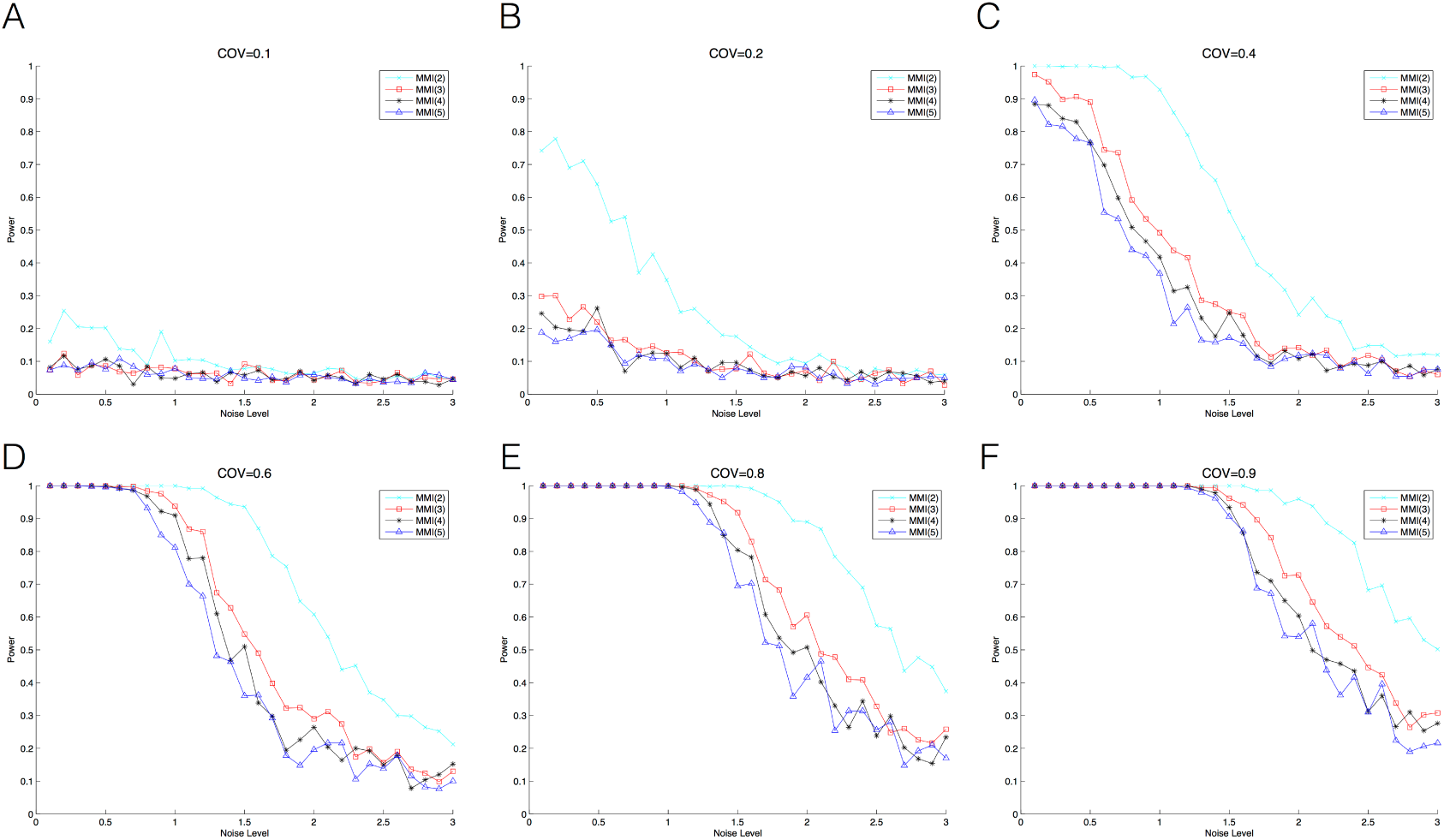
Power and noise tolerance for MMI by assigning different number of modes. The numbers in brackets denote the number of modes. **(A)** Covariance=0.1, **(B)** Covariance=0.2, **(C)** Covariance=0.4, **(D)** Covariance=0.6, **(E)** Covariance=0.8, **(F)** Covariance=0.9.

### 3.3 Simulation results for MDI

We simulate five tree structures, respectively of 10, 20, 50, 100 and 200 nodes. For each tree structure, we define a spectrum of covariances from weak to strong (0.1, 0.2, 0.4, 0.6 and 0.8), as well as a uniformly distributed noise. When noise is neglected, MDI consistently performs the best (Table 1 (a), 2 (a), 3 (a), 4 (a), 5 (a)). This advantage is very significant in the case of small covariance (0.1) with large number of nodes (200), where the AUC of MDI is 15.7% higher than the second best method, that is, CLR. MDI performs the best in 69 out of 75 simulations (92%). The 6 cases where MDI does not achieve the best result are the cases of high noise level, in which ARACNE or CLR show better performance (Table 1 (c), 2 (c), 5 (b), 5 (c)). However, the AUC values of MDI remain comparable in these cases. We note that the three MI-based methods, MDI, ARACNE and CLR, to perform better than MaxEnt, GENIE3 and TIGRESS in general, which suggests that MI-based methods may be more appropriate for capturing regulatory relationship.

**Table 1.** Simulation result for 10 nodes tree by comparing MDI with other methods in terms of AUC.

| (a) Simulation without any noise |  |  |  |  |  |  |
| --- | --- | --- | --- | --- | --- | --- |
|  | ARACNE | CLR | GENIE3 | MaxEnt | TIGRESS | MDI |
| 0.1 | 0.64 | 0.79 | 0.57 | 0.58 | 0.53 | **0.81** |
| 0.2 | 0.92 | 0.97 | 0.68 | 0.73 | 0.60 | **1** |
| 0.4 | **1** | **1** | 0.89 | 0.76 | 0.71 | **1** |
| 0.6 | **1** | **1** | 0.92 | 0.47 | 0.56 | **1** |
| 0.8 | **1** | **1** | 0.84 | 0.68 | 0.55 | **1** |
| (b) Simulation with 3/5 random noise |  |  |  |  |  |  |
|  | ARACNE | CLR | GENIE3 | MaxEnt | TIGRESS | MDI |
| 0.1 | 0.64 | 0.64 | 0.59 | 0.61 | 0.53 | **0.75** |
| 0.2 | 0.88 | 0.88 | 0.54 | 0.70 | 0.58 | **1** |
| 0.4 | 0.96 | **1** | 0.72 | 0.77 | 0.7289 | **1** |
| 0.6 | 0.89 | **1** | 0.83 | 0.49 | 0.5643 | **1** |
| 0.8 | **1** | **1** | 0.66 | 0.6543 | 0.5643 | **1** |
| (c) Simulation with 6/5 random noise |  |  |  |  |  |  |
|  | ARACNE | CLR | GENIE3 | MaxEnt | TIGRESS | MDI |
| 0.1 | **0.63** | 0.40 | 0.46 | 0.61 | 0.56 | 0.46 |
| 0.2 | 0.60 | 0.71 | 0.50 | 0.67 | 0.54 | **0.87** |
| 0.4 | 0.85 | **1** | 0.80 | 0.58 | 0.65 | **1** |
| 0.6 | 0.79 | **0.99** | 0.71 | 0.53 | 0.57 | **0.99** |
| 0.8 | 0.94 | 0.97 | 0.73 | 0.65 | 0.55 | **0.98** |

**Table 2.** Simulation result for 20 nodes tree by comparing MDI with other methods in terms of AUC.

| (a) Simulation without any noise |  |  |  |  |  |  |
| --- | --- | --- | --- | --- | --- | --- |
|  | ARACNE | CLR | GENIE3 | MaxEnt | TIGRESS | MDI |
| 0.1 | 0.69 | 0.74 | 0.4541 | 0.47 | 0.49 | **0.87** |
| 0.2 | 0.89 | 0.98 | 0.52 | 0.42 | 0.44 | **1** |
| 0.4 | **1** | **1** | 0.87 | 0.61 | 0.65 | **1** |
| 0.6 | **1** | **1** | 0.90 | 0.57 | 0.54 | **1** |
| 0.8 | **1** | **1** | 0.93 | 0.64 | 0.61 | **1** |
| (b) Simulation with 3/5 random noise |  |  |  |  |  |  |
|  | ARACNE | CLR | GENIE3 | MaxEnt | TIGRESS | MDI |
| 0.1 | 0.52 | 0.53 | 0.35 | 0.47 | 0.50 | **0.73** |
| 0.2 | 0.87 | 0.93 | 0.56 | 0.43 | 0.45 | **1** |
| 0.4 | **1** | **1** | 0.77 | 0.61 | 0.62 | **1** |
| 0.6 | **1** | **1** | 0.83 | 0.55 | 0.54 | **1** |
| 0.8 | 0.98 | **1** | 0.85 | 0.60 | 0.59 | **1** |
| (c) Simulation with 6/5 random noise |  |  |  |  |  |  |
|  | ARACNE | CLR | GENIE3 | MaxEnt | TIGRESS | MDI |
| 0.1 | 0.53 | 0.61 | 0.47 | 0.50 | 0.49 | **0.69** |
| 0.2 | 0.59 | 0.67 | 0.5324 | 0.50 | 0.48 | **0.86** |
| 0.4 | 0.78 | 0.87 | 0.6340 | 0.61 | 0.66 | **0.99** |
| 0.6 | 0.97 | **0.99** | 0.65 | 0.49 | 0.47 | **0.99** |
| 0.8 | 0.88 | **0.99** | 0.75 | 0.64 | 0.57 | 0.98 |

**Table 3.** Simulation result for 50 nodes tree by comparing MDI with other methods in terms of AUC.

(a) Simulation without any noise
|  | ARACNE | CLR | GENIE3 | MaxEnt | TIGRESS | MDI |
| --- | --- | --- | --- | --- | --- | --- |
| 0.1 | 0.53 | 0.79 | 0.54 | 0.49 | 0.53 | **0.94** |
| 0.2 | 0.93 | **1** | 0.62 | 0.52 | 0.53 | **1** |
| 0.4 | **1** | **1** | 0.85 | 0.61 | 0.56 | **1** |
| 0.6 | **1** | **1** | 0.95 | 0.61 | 0.57 | **1** |
| 0.8 | **1** | **1** | 0.95 | 0.56 | 0.57 | **1** |

(b) Simulation with 3/5 random noise
|  | ARACNE | CLR | GENIE3 | MaxEnt | TIGRESS | MDI |
| --- | --- | --- | --- | --- | --- | --- |
| 0.1 | 0.50 | 0.63 | 0.50 | 0.48 | 0.50 | **0.83** |
| 0.2 | 0.80 | 0.94 | 0.60 | 0.5279 | 0.51 | **1** |
| 0.4 | 0.99 | **1** | 0.78 | 0.63 | 0.58 | **1** |
| 0.6 | **1** | **1** | 0.88 | 0.62 | 0.60 | **1** |
| 0.8 | **1** | 0.99 | 0.87 | 0.55 | 0.54 | **1** |

(c) Simulation with 6/5 random noise
|  | ARACNE | CLR | GENIE3 | MaxEnt | TIGRESS | MDI |
| --- | --- | --- | --- | --- | --- | --- |
| 0.1 | 0.48 | 0.52 | 0.51 | 0.46 | 0.50 | **0.64** |
| 0.2 | 0.55 | 0.64 | 0.49 | 0.47 | 0.50 | **0.86** |
| 0.4 | 0.82 | 0.90 | 0.64 | 0.57 | 0.55 | **1** |
| 0.6 | 0.96 | 0.99 | 0.75 | 0.60 | 0.57 | **1** |
| 0.8 | 0.97 | 0.99 | 0.84 | 0.60 | 0.59 | **1** |

**Table 4.** Simulation result for 100 nodes tree by comparing MDI with other methods in terms of AUC.

(a) Simulation without any noise
|  | ARACNE | CLR | GENIE3 | MaxEnt | TIGRESS | MDI |
| --- | --- | --- | --- | --- | --- | --- |
| 0.1 | 0.66 | 0.74 | 0.54 | 0.53 | 0.54 | **0.91** |
| 0.2 | 0.90 | **1** | 0.64 | 0.60 | 0.54 | **1** |
| 0.4 | **1** | **1** | 0.82 | 0.54 | 0.54 | **1** |
| 0.6 | **1** | **1** | 0.96 | 0.60 | 0.58 | **1** |
| 0.8 | **1** | **1** | 0.97 | 0.56 | 0.55 | **1** |

(b) Simulation with 3/5 random noise
|  | ARACNE | CLR | GENIE3 | MaxEnt | TIGRESS | MDI |
| --- | --- | --- | --- | --- | --- | --- |
| 0.1 | 0.56 | 0.66 | 0.56 | 0.52 | 0.52 | **0.80** |
| 0.2 | 0.86 | 0.92 | 0.60 | 0.58 | 0.52 | **0.98** |
| 0.4 | **1** | **1** | 0.74 | 0.54 | 0.53 | **1** |
| 0.6 | **1** | **1** | 0.92 | 0.60 | 0.57 | **1** |
| 0.8 | **1** | **1** | 0.93 | 0.54 | 0.56 | **1** |

(c) Simulation with 6/5 random noise
|  | ARACNE | CLR | GENIE3 | MaxEnt | TIGRESS | MDI |
| --- | --- | --- | --- | --- | --- | --- |
| 0.1 | 0.62 | 0.56 | 0.55 | 0.52 | 0.52 | **0.63** |
| 0.2 | 0.63 | 0.65 | 0.56 | 0.57 | 0.53 | **0.82** |
| 0.4 | 0.76 | 0.93 | 0.63 | 0.52 | 0.51 | **0.99** |
| 0.6 | 0.97 | **1** | 0.79 | 0.58 | 0.56 | **1** |
| 0.8 | 0.96 | **1** | 0.81 | 0.56 | 0.57 | 0.99 |

**Table 5.** Simulation result for 200 nodes tree by comparing MDI with other methods in terms of AUC.

(a) Simulation without any noise
|  | ARACNE | CLR | GENIE3 | MaxEnt | TIGRESS | MDI |
| --- | --- | --- | --- | --- | --- | --- |
| 0.1 | 0.6432 | 0.7172 | 0.5277 | 0.49 | 0.50 | **0.83** |
| 0.2 | 0.8780 | 0.9732 | 0.6040 | 0.51 | 0.49 | **1** |
| 0.4 | **1** | **1** | 0.84 | 0.59 | 0.57 | **1** |
| 0.6 | **1** | **1** | 0.97 | 0.61 | 0.62 | **1** |
| 0.8 | **1** | **1** | 0.98 | 0.59 | 0.63 | **1** |

(b) Simulation with 3/5 random noise
|  | ARACNE | CLR | GENIE3 | MaxEnt | TIGRESS | MDI |
| --- | --- | --- | --- | --- | --- | --- |
| 0.1 | 0.56 | 0.63 | 0.53 | 0.4901 | 0.51 | **0.75** |
| 0.2 | 0.71 | 0.89 | 0.56 | 0.52 | 0.51 | **0.97** |
| 0.4 | 0.99 | **1** | 0.75 | 0.60 | 0.56 | **1** |
| 0.6 | **1** | 0.91 | 0.98 | 0.61 | 0.60 | **1** |
| 0.8 | **1** | **1** | 0.94 | 0.59 | 0.63 | 0.96 |

(c) Simulation with 6/5 random noise
|  | ARACNE | CLR | GENIE3 | MaxEnt | TIGRESS | MDI |
| --- | --- | --- | --- | --- | --- | --- |
| 0.1 | 0.50 | **0.51** | 0.49 | 0.48 | 0.49 | 0.49 |
| 0.2 | 0.54 | 0.52 | 0.51 | 0.53 | 0.49 | **0.59** |
| 0.4 | 0.58 | 0.66 | 0.61 | 0.54 | 0.55 | **0.74** |
| 0.6 | 0.68 | 0.78 | 0.66 | 0.58 | 0.60 | **0.86** |
| 0.8 | 0.83 | 0.90 | 0.72 | 0.58 | 0.61 | **0.90** |

### 3.4 Transcriptome components analysis for large expression compendium

Recently, Fehrmann *et al.* Fehrmann et al. (2015) propose a new perspective that the transcriptome components, as an underlying regulatory factor, can influence a batch of target gene expressions. The transcriptome components are calculated by principal component analysis on a correlation matrix, each of them capture a proportion of variance in the correlation space. The correlation matrix used in their study is computed with Pearson correlation. In this experiment, we compute a correlation matrix with MMI, and examine if the matrix could result in similar, or better results.

We evaluate the transcriptome components with Cronbach’s *α* value; transcriptome components with the values larger than 0.7 are generally considered as high quality ones and can be used to predict gene functions in the future. We further evaluate the biological function enrichment of principal components by grouping genes’ coefficients according to MSigDB Subramanian et al. (2005). The gene set enrichment *Z*-score is computed by comparing the coefficients between the genes belonging and not belonging to the particular function by two-sample *t*-test. These TCs are regarded as enriched in a function term if the *p*-values are less than 0.05, after 10,000 simulations.

The results are as shown in Figure fig:Figure4. Using the matrix obtained from MMI, we find 657 transcriptome components with high quality, in which 650 transcriptome components are enriched in at least one biological function in MSigDB. On the other hand, for the matrix obtained from Pearson correlation, the corresponding values are only 379 and 369. Thus, the use of MMI achieves nearly twice functional TCs than Pearson correlation. These functional transcriptome components can be further used to predict potential gene functions.

**Figure 4.**
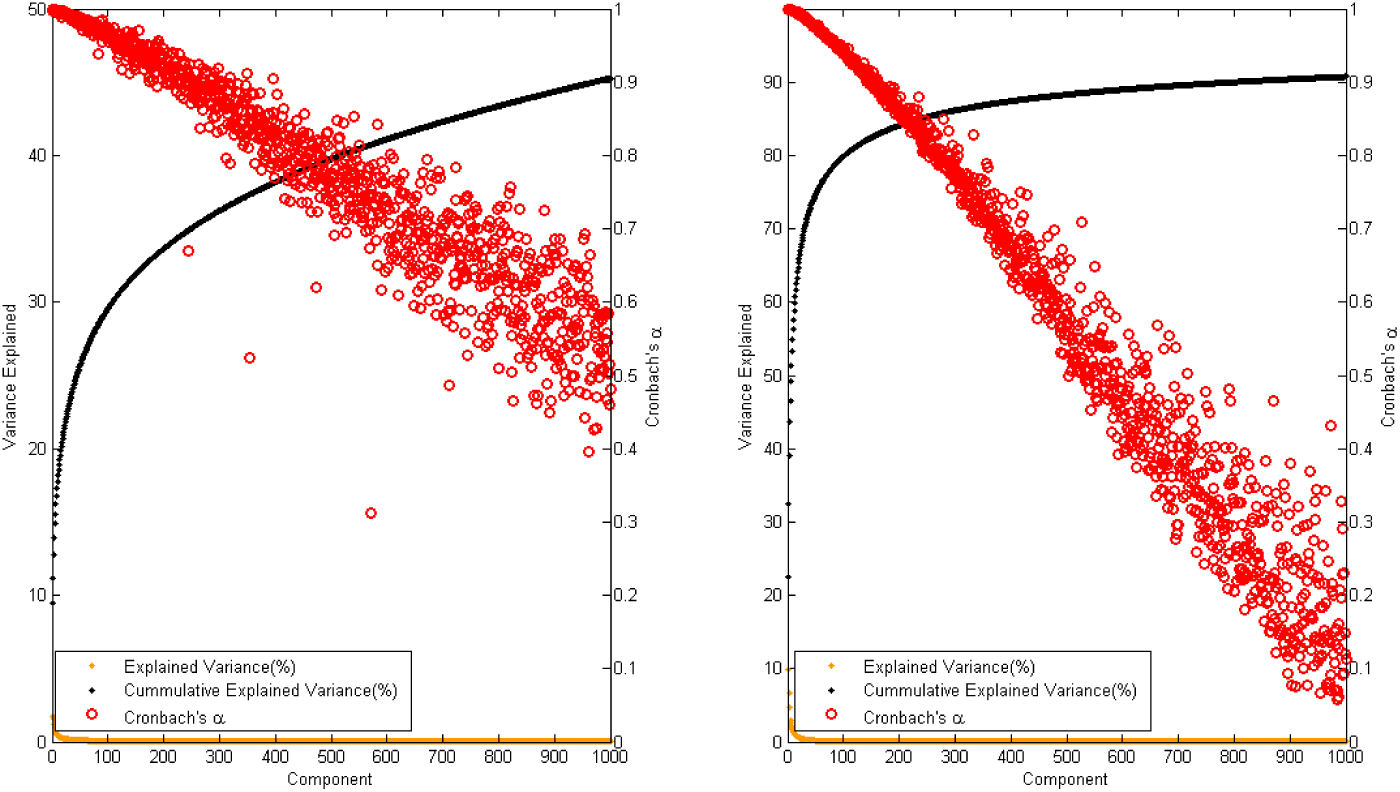
Covariance explained by eigenvalues and their Cronbach’s *α* values. (**Left**: MMI, **Right**: Pearson correlation). We choose top 1000 principle components to calculated the variance they explained and their stability.

### 3.5 Apply to the exome sequencing for familial Mendelian disorders

Exome sequencing has been widely applied in identifying disease causal genes in the families affected by rare Mendelian disorders. However, it is a formidable task to pinpoint the causal one from very large amount of candidate genes. Erlich *et al.* Erlich et al. (2011) propose a novel method to remove meaningless genes by disease network analysis, working under the assumption that the disease causal gene should share biological functions with the other established genes of the disease. After bioinformatics analysis, there remains 15 candidate genes to be determined. Instead of utilizing interactions from databases such as Gene Ontology, KEGG pathway, we propose to evaluate gene-gene interactions by their co-expressions. We remove five genes that are missing in the Affymetrix HG U133A and calculate the average co-expression for each candidate gene with the ten established genes (**Supplementary note**) associated with HSP by MMI. Among the 10 candidate genes, the causal gene *KIF1A* is the only one significantly co-expresses with established genes (*FDR <* 0.05) after 10,000 simulations (Figure fig:Figure5).

**Figure 5.**
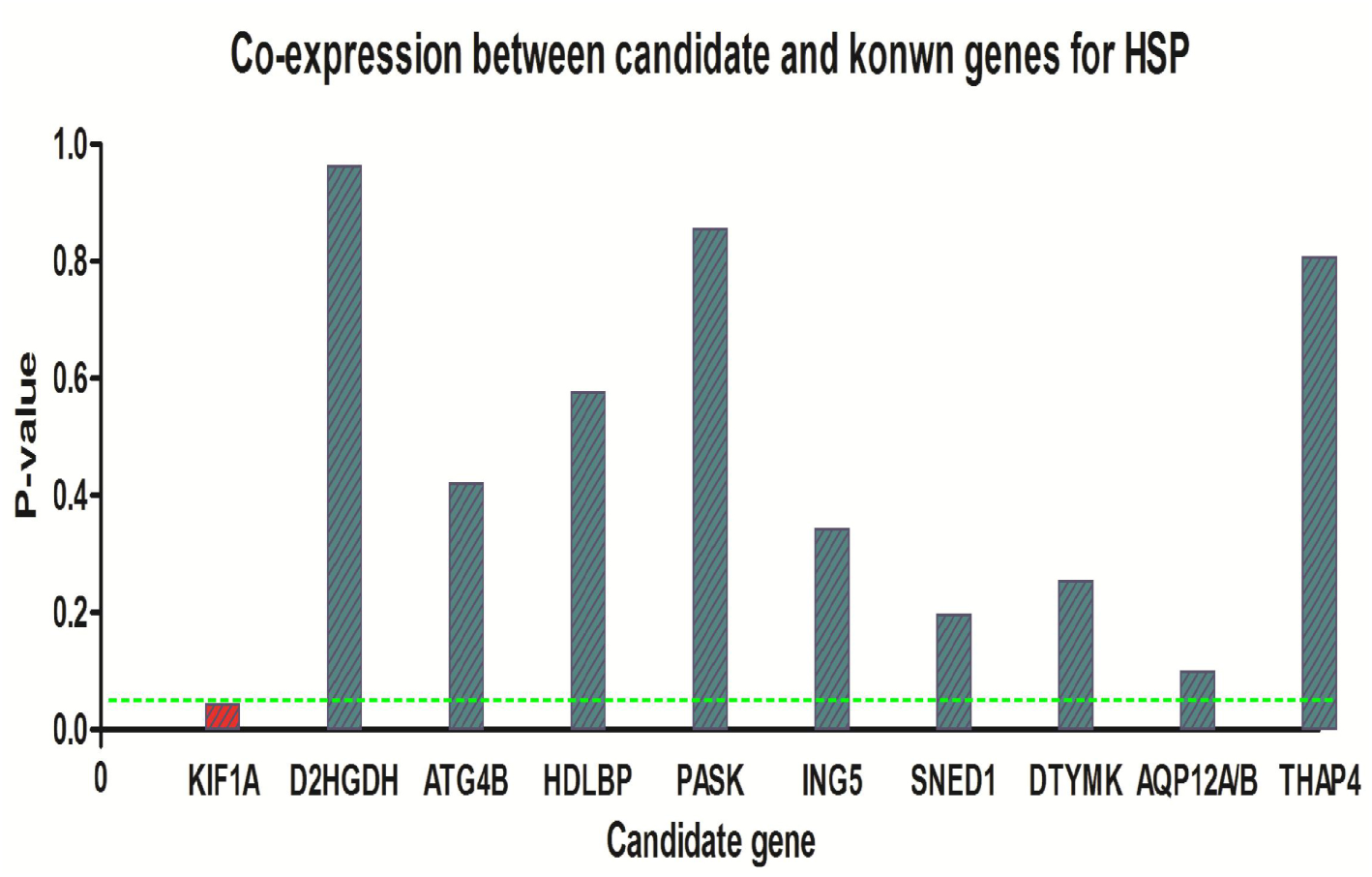
The P-values calculated by their average co-expression with known genes of pure HSP after 10,000 times simulation. The co-expression values are calculated by MMI. KIF1A, the real disease causal gene, is the only significant one comparing with other 10 candidate genes. The green dashed line represents the P-value=0.05.

### 3.6 Apply to *de novo* mutations to identify the ‘hot gene’

*De novo* mutations have been proven to be associated with many neurodevelopment disorders Iossifov et al. (2014); Fromer et al. (2014), especially for schizophrenia and ASDs. These genes with *de novo* mutations do not act individually, they are enriched in the same protein-protein interactions, gene ontology term or regulatory network De Rubeis et al. (2014). We apply MDI to identify the ‘hot gene’, which organize the other genes in the network. We construct the directly interaction network by involving 33 candidate genes (**Supplementary note**) identified in the latest family based study of ASDs De Rubeis et al. (2014) (with TADA He et al. (2013) *FDR <* 0.1). MDI identifies *ANK2* as the ‘hot gene’, with the most connections with the other genes (Figure fig:Figure6).

**Figure 6.**
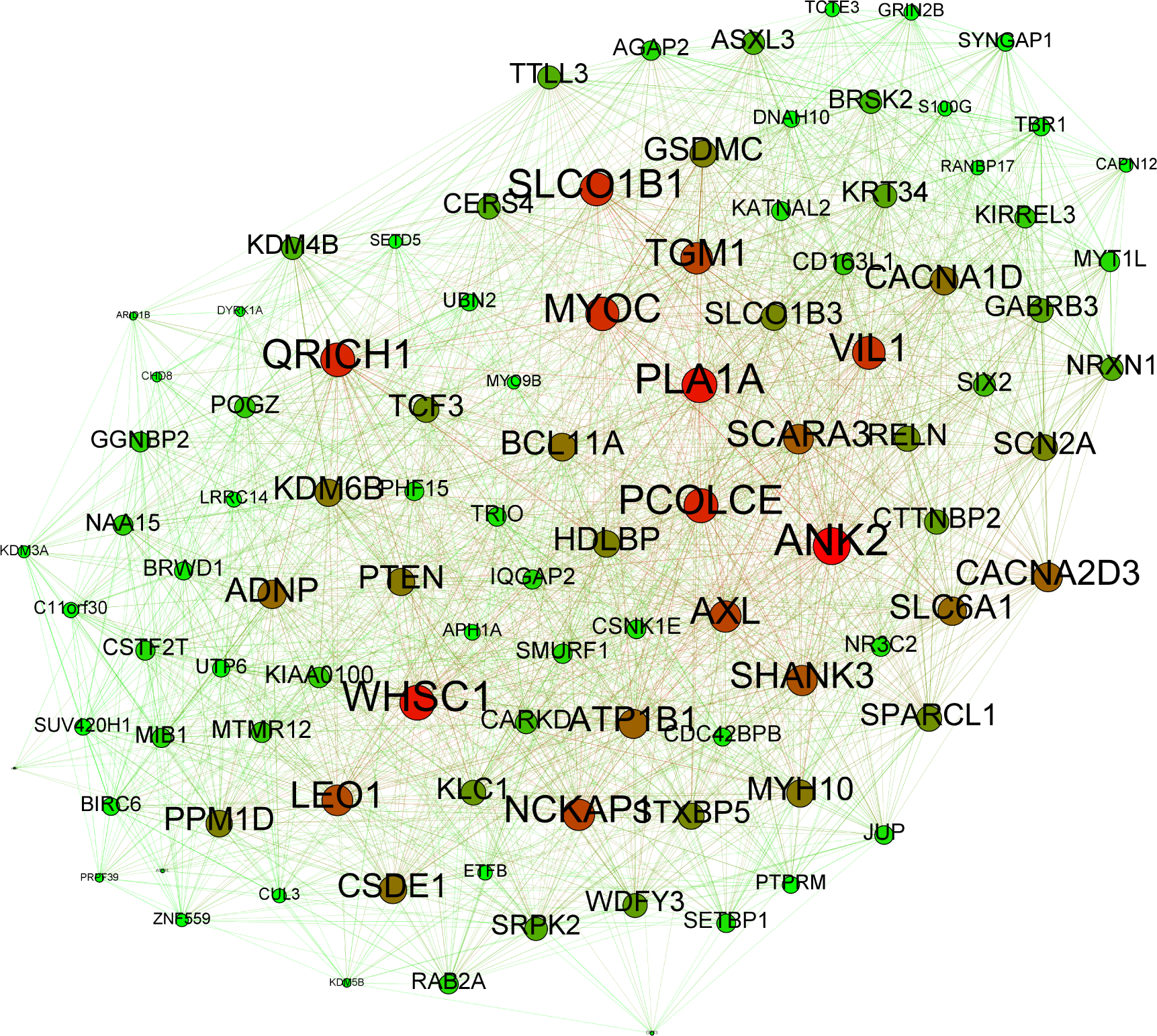
The ‘hot gene’ calculated by the weighted connective degree for each candidate genes. The size and color of nodes represent their weighted connective degree and expression level, respectively.

*ANK2* has been detected to contain recurrent *de novo* mutations recently Willsey et al. (2013), and is one of the five ‘high-confidence’ ASDs candidate genes (the other four are *CHD8, DYRK1A, GRIN2B, SCN2A*). The expression of *ANK2* peaks slightly during mid-fetal development, which is the crucial time for ASDs risk. *ANK2*’s expression closely matches that of many other ASDs candidate genes, including *SCN2A* Willsey et al. (2013). In 1991, a study conducted by Kordeli and Bennett Kordeli and Bennett (1991) finds that knockout mice’s *ANK2* may lead to their lack of brain structures called the corpus callosum—a symptom which is likely related to ASDs; about one-third of the patients with corpus callosum are also affected by ASDs. These evidences strongly suggests to us that MDI may have identified the key gene in ASDs; *ANK2* may play a key role in ASDs pathogenesis.

### 3.7 MicroRNA targets prediction

We have noticed that, in HOCTAR, the microRNA targets is predicted by jointly considering the sequence feature and expression correlation between targets and microRNA host gene. But it may produce false targets due to their functional similarity with the real targets. This may result in their co-expression with each other. In HOCTAR, the Pearson correlation can barely distinguish transitive co-expression and direct regulation. In this test, we do a comparison between MDI and Pearson correlation used in HOCTAR as well as other three sequence-based prediction software-TargetScan Lewis et al. (2005), miRnada John et al. (2004) and PicTar Krek et al. (2005). To remove the false positives, we extract the putative targets that are confirmed by at least two of the three sequence-based software mentioned. The intragenic microRNAs and their host genes are derived from miRIAD Malone et al. (2013). The validated microRNA-target pairs used as benchmark are collected by literature search (**TableS2**). In Figure fig:Figure7, we illustrate the rank of validated microRNA-target pairs for each microRNA against the percentile. For MDI, there are 90.70% (78/86) of the validated pairs locate in the top 50 percentile, which outperform Pearson correlation (63.95%,55/86). The three sequence-based prediction algorithms perform well, TargetScan 69.33% (52/75), miRanda 78.87% (56/71) and PicTar 74.19% (23/31). This suggests that acceptable prediction can already be achieved using only the overlap of sequence-based prediction software. Incorporating expression correlation can increase the accuracy of microRNA target prediction, but it is important to remove the transitive correlations. (In Figure fig:Figure8 we extract the reliable targets with greater MDI than the validated targets for each microRNA.)

**Figure 7.**
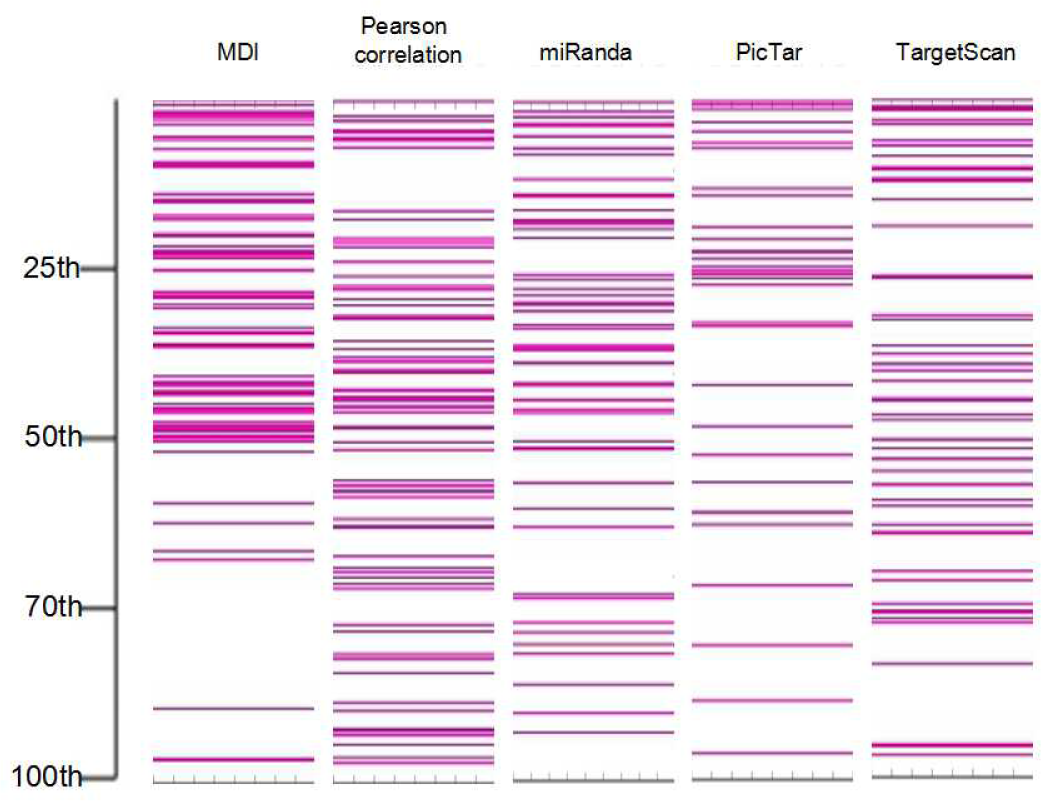
The performance of MDI to recognize previous validated microRNA targets. MDI is compared with Pearson correlation and three sequence based target prediction approaches (miRanda, PicTar, TargetScan), respectively. The ranks of validated targets (pink line) are demonstrated as the percentile among the all predicted results for each microRNA.

**Figure 8.**
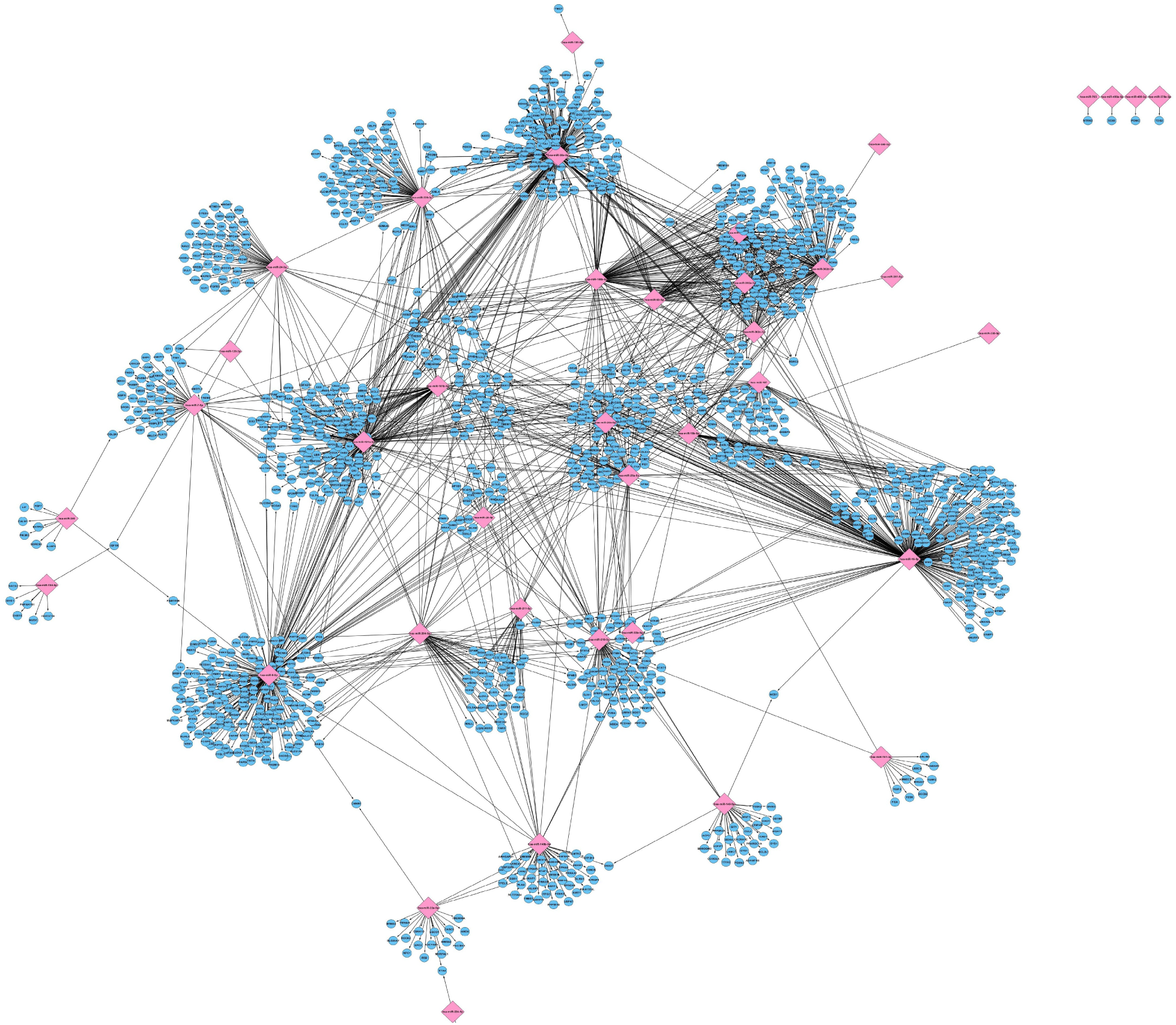
The microRNA regulatory network predicted by MDI. For each microRNA, we plot all the predicted targets with greater MDI than the validated ones. The pink diamonds and blue circles present microRNAs and their targets, respectively.

## 4 DISCUSSION

Bioinformatics data analysis task always encounters sample size issue, especially for biomedical data, which suffer from pathogenic difference and sample heterogeneity. While the cost for high-throughput technology has decreased dramatically in recent years, the collection of extremely large samples is still luxury that few studies can afford. Integrating the huge amount of biological data readily available from public databases remains the best option. Several studies has proved the superiority of data integration, much attention has been paid on how to efficiently store and search useful knowledge from large expression compendium. However, the integration of these database is plagued by pathogenic difference and sample heterogeneity.

The genes do not act alone, they tend to be grouped together and with particular topological structure, we call it as pathway. The pathway relies on the interactions between genes, including causal and non-causal interactions. The causal interactions refer to regulatory interactions, the regulators or their products physically bind to those target genes’ sequences to make their status changed. The non-causal interactions are a kind of indirect interactions, where the genes interact with each other through transitive ones. Intuitively, these genes may share similar functions. Several databases have collected many experimental validated interactions. But such data are usually incomplete and biased, for example only a small proportion of transcription factors targets are well studied. We require an approach that can explore the interactions from unstructured data based on their inherent activities.

Such large amount of gene expression data shed light on evaluating the gene-gene interactions, which can be detected by computing expression dependencies. On the other hand, it is an uneasy task because traditional methods are no longer appropriate for large expression collection. Previous findings suggest that gene expressions from multiple datasets follow a Gaussian mixture models. Inspired by this observation, we propose MMF, a framework that depicts the gene expression data by Gaussian mixtures. Each mode captures one type of “local features”, which denotes noise or particular cellular status, hence allowing the heterogeneous data sets to be integrated naturally. Two measures, MMI and MDI, are defined over the framework to capture gene-gene co-expression and regulatory interactions, respectively. They outperform other measures in the simulation tests, and several real data benchmarks proved their practicality to detect important interactions or genes.

MIC is a comparable method that can detect novel associations in large datasets Reshef et al. (2011). It also considers the “local feature” to improve the accuracy and resist the noisy influence. Two major drawbacks prohibit MIC to be widely applied in large expression data. First, as an improvement of B-spline, MIC considers the best grid partition to calculate mutual information for gene pairs. But from our simulations, usually MIC cannot find the best choice for the data from Gaussian mixture models. In addition, MIC ignores the expression correlation within each grid, which makes the power decreased; second, MIC attempts the partition for every pair of genes, which introduce a huge computational burden. MMI explicitly solves these two problems and achieves better performance.

In the future, we plan to apply MMF to other types of continuous data and extend it to other underlying distributions such as Poisson distribution or Negative binomial distribution. We believe MMF can find wide application in discovering new interactions from integrated gene expression data, as well as help in furthering the analysis of big biomedical data.

## 5 CONCLUSION

The fast accumulation of high-throughput gene expression data provides us an unprecedented opportunity to understand the gene-gene interactions and prioritize the disease candidate genes. However, most of the previous approaches can not accurately depict gene expression profiles from large expression compendium due to considerable noise and heterogeneity between samples. We propose a new statistical measure Multimodal framework to model gene expressions with mixtures of Gaussian distributions, which is further extended to Multimodal Mutual information and Multimodal Direct information for calculating gene-gene co-expression and gene regulation, respectively. The practical use of MMF is further demonstrated in three biological applications: 1. Prioritizing KIF1A as the candidate causal gene of HSP from familial exome sequencing data; 2. Detecting ANK2 as the ‘hot genes’ for ASDs, derived from exome sequencing family based study; 3. Predicting the microRNA target genes based on both sequence and expression information. We believe MMF can be served as a general framework for discovering relationships within very massive biomedical datasets.

## AUTHOR CONTRIBUTIONS

SL supervised the work and together with LZ, developed MMF and procedure of experiment. LZ implemented the MMI method in matlab. LZ, JC did the experiments on simulation and real data. LZ and SL wrote the manuscript. All authors have read and approved the final manuscript.

## ACKNOWLEDGMENTS

We would like to thank Yen Kaow Ng for informative discussions.

## SUPPLEMENTAL DATA

Supplementary Material The Supplementary Material for this article can be found online at: XXX

