## Supplementary Note for "A multimodal framework for detecting direct and indirect gene-gene interactions from large expression compendium"

### Supplementary Material

#### Maximum entropy

Suppose the expression level for $G$ genes can be denoted as a vector $\vec{x} = (x_1, ..., x_g)$, the number of cluster each gene sampled from is $C = (c_1, ..., c_g)$, the sample T are demonstrated as $\vec{x}^1 ... \vec{x}^N$. We can translate the dependence among the genes into Kullback-Leibler divergence between the joint probability density (JPD) function and the product of the marginal distributions,

$$I[P] = \sum_{i=1}^{G} P(\vec{x}) log_2 \frac{P(\vec{x})}{\prod_i P(x_{i,k_1})} = \langle log_2 \frac{P(\vec{x})}{\prod_i P(x_{i,k_1})} \rangle = \sum_i S[X_{i,k_1}] - S[\mathbf{X}] \tag{S1}$$

Assuming u-ways marginals are known, our aim is to found approximation of JPD, $P(x_{i,k_1}, x_{j,k_2}, ..., x_{u,k_u})$ ,only respects the marginals, without any additional assumptions about the JPD.

$$\begin{aligned} P(\vec{x}) = \frac{1}{Z} \exp[-\sum_{i}^{G} \varphi_i(x_{i,k}) - \sum_{i,j}^{G} \varphi_{ij}(x_{i,k_1}, x_{j,k_2}) \\ - \sum_{i,j,u}^{G} \varphi_{iju}(x_{i,k_1}, x_{j,k_2}, x_{u,k_u}) \\ \equiv e^{-H\{g_{i,k_1}\}} \end{aligned} \tag{S2}$$

Z is the normalization factor, $\varphi$ is called potentials, $-H(\{g_{i,k_1}\})$ is Hamilton function. Within such a model, we assert that a set of variables interact if and only if the high-order potentials that depend exclusively on these variables are nonzero. A reasonable way to specify $\varphi$ based on maximum entropy approximations to u-ways interaction. Because the limited number of sample size, we only consider the potential of two-way interaction to maximize

$$S = -\sum_{\vec{x}} p(\vec{x}) lnp(\vec{x}) \tag{S3}$$

subject to the constrains

$$\sum_{\vec{x}} p(\vec{x}) = 1 \tag{S4}$$

$$\langle x_{i,k_1} \rangle = \sum_{\vec{x}} x_{i,k_1} = \frac{1}{N} \sum_{t=1}^{N} (x_{i,k_1})^t \tag{S5}$$

$$\langle x_{i,k_1} x_{j,k_2} \rangle = \sum_{\vec{x}} x_{i,k_1} x_{j,k_2} = \frac{1}{N} \sum_{t=1}^{N} (x_{i,k_1})^t (x_{j,k_2})^t \tag{S6}$$

Eq.4 normalize the observed probability of genes sum to 1. Eq.5 and 6 preserve the mean expression and two-way interactions between models. On the basis of these constrains, we can refine the covariance matrix $Cov$

$$Cov_{x_{i,k_1},x_{j,k_2}} = \langle x_{i,k_1} x_{j,k_2} \rangle - \langle x_{i,k_1} \rangle \langle x_{j,k_2} \rangle \tag{S7}$$

$$\varsigma = S - \nu \sum_{\vec{x}} - \sum_{i=1}^{G} \mu_i \sum_{\vec{x}} p(\vec{x}) x_{i,k_1} - \sum_{i,j=1}^{G} \lambda_{ij} \sum_{\vec{x}} p(\vec{x}) x_{i,k_1} x_{j,k_2} \tag{S8}$$

let $\frac{\partial \varsigma}{\partial p(\vec{x})} = 0$ and $M \equiv \frac{1}{2}\lambda$, we have

$$p(\vec{x}) = e^{-1-\nu-\vec{\mu}\cdot\vec{x}-\frac{1}{2}\vec{x}M\vec{x}} = Ae^{-\frac{1}{2}\vec{y}M\vec{y}} \tag{S9}$$

where $\vec{y} = \vec{x} + \vec{\mu}M^{-1}$ and $A = e^{\frac{1}{2}\vec{\mu}M^{-1}\vec{\mu}} e^{-1-\nu}$. By simply assuming the continuous distribution of transcripts in the genome, the summation of Eq.4-6 can be replaced by integral. Then the Eq.4-6 can be transformed to

$$1 = \int d^G x p(\vec{x}) = \int d^G y A e^{-\frac{1}{2}\vec{y}M\vec{y}} \tag{S10}$$

$$\begin{aligned} \langle x_{i,k_1} \rangle &= \int d^G x p(\vec{x}) x_{i,k_1} \\ &= \int d^G y A e^{-\frac{1}{2}\vec{y}M\vec{y}} (y_{i,k_1} - \sum_j (M_{i,j,k_1,k_2})^{-1} \mu_j) \\ &= -\sum_j (M_{i,j,k_1,k_2})^{-1} \mu_{j,k_2} \end{aligned} \tag{S11}$$

$$\begin{aligned} &\langle x_{i,k_1} x_{j,k_2} \rangle \\ &= \int d^G x p(\vec{x}) x_{i,k_1} x_{j,k_2} \\ &= \int d^G y A e^{-\frac{1}{2}\vec{y}M\vec{y}} [\langle x_{i,k_1} \rangle \langle x_{j,k_2} \rangle + y_{i,k_1} \langle x_{j,k_2} \rangle + y_{j,k_2} \langle x_{i,k_1} \rangle + y_{i,k_1} y_{j,k_2}] \\ &= \langle x_{i,k_1} \rangle \langle x_{j,k_2} \rangle + \int A e^{-\frac{1}{2}\vec{y}M\vec{y}} y_i y_j \end{aligned} \tag{S12}$$

To solve the integral in Eq.12, we can define a generating function

$$Z(\vec{J}) = \int d^G y e^{\frac{1}{2}\vec{y}M\vec{y} + \vec{J}\cdot\vec{y}} \tag{S13}$$

By making the substitutions $\vec{z} = \vec{y} - M^{-1}\vec{J}$, the Eq.13 can be re-written as

$$Z(\vec{J}) = e^{\frac{1}{2}\vec{J}M^{-1}\vec{J}} \int d^G z e^{-\frac{1}{2}\vec{z}} = \frac{(2\pi)^{\frac{N}{2}}}{\sqrt{detM}} e^{\frac{1}{2}\vec{J}M^{-1}\vec{J}} \tag{S14}$$

The derivation of Eq.14 can yield the pairwise correlation

$$\langle y_{i,k_1} y_{j,k_2} \rangle = \frac{1}{Z(\vec{J})} \frac{\delta^2}{\delta J_{i,k_1} \delta J_{j,k_2}} Z(\vec{J})|_{\vec{J}=0} = (M)^{-1}_{i,j,k_1,k_2} \tag{S15}$$

combining with Eq.12, we find that the interaction matrix is simply the inverse of covariance matrix.

### Candidate and known gene list for HSP

HSP candidate genes: KIF1A,D2HGDH,ATG4B,HDLBP,PASK,ING5,SNED1, DTYMK,AQP12A/B,THAP4
HSP known causal genes: CYP7B1,HSPD1,KIAA0196,KIFSA,NIPA1,PLP1,PNPLA6, REEP1,SPAST,ATL1,ZFYVE27

### Candidate gene list for ASDs

The candidate genes list for identifying hot gene of ASDs are: SCN2A,SYNGAP1,CHD8,ARID1B,ANK2, SUV420H1,DYRK1A,GRIN2B,ADNP,TBR1,POGZ,CUL3, KATNAL2,BCL11A,CACNA2D3,MIB1, GABRB3,CTTNBP2,MLL3,PTEN,RELN,ASXL3,MYO9B,ETFB,ASH1L,TRIO,NAA15,MYT1L,NR3C2, APH1A,SETD5,VIL1 CDC42BPB
